# Following the p63/Keratin5 Basal Cells in the Sensory and Non-sensory Epithelia of The Vomeronasal Organ

**DOI:** 10.1101/2023.11.16.567453

**Authors:** Noah M. LeFever, Raghu Ram Katreddi, Nikki M. Dolphin, Nick A. Mathias, Paolo E. Forni

## Abstract

The Vomeronasal organ (VNO) is a part of the accessory olfactory system, which is responsible for detecting pheromones, chemical factors that trigger a spectrum of sexual and social behaviors. The vomeronasal epithelium (VNE) shares several features with the epithelium of the main olfactory epithelium (MOE). However, it is a distinct neuroepithelium populated by chemosensory neurons that differ from the olfactory sensory neurons (OSNs) in cellular structure, receptor expression, and connectivity.

The vomeronasal organ of rodents comprises a sensory epithelium and a thin non-sensory epithelium that morphologically resembles the respiratory epithelium.

Sox2-positive cells have been previously identified as the stem cell population that gives rise to neuronal progenitors in MOE and VNE. In addition to these, the MOE also comprises p63 positive horizontal basal cells (HBCs), a second pool of quiescent stem cells that become active in response to injury.

Immunolabeling against the transcription factor p63, Keratin-5 (Krt5), Krt14 and Krt5Cre tracing experiments highlighted the existence of horizontal basal cells distributed along the basal lamina of the VNO forming from progenitors along the basal lamina of the marginal zones. Moreover, these experiments revealed that the NSE of rodents is, like the respiratory epithelium, a stratified epithelium where the p63/Krt5+ basal cells self-replicate and give rise to the apical columnar cells facing the lumen of the VNO.

## Introduction

Stem cells are undifferentiated cells that have the ability to self-renew or differentiate into diverse cell types (Weissman, 2000). Adult tissues can contain stem cells that stay mitotically active to maintain tissue homeostasis or in a quiescent state (Li and Clevers, 2010). Interestingly, the quiescent stem cell population can become active in response to extrinsic cell signals from the surrounding niche or stress signals post-injury (Cheung and Rando, 2013; Fletcher *et al*., 2011; Hsu and Fuchs, 2012).

The mouse olfactory system is a well-characterized neuronal system in its ability to turn over new chemosensory neurons throughout life and to regenerate the sensory epithelium during post-injury conditions (Schwob *et al*., 2017). In mice, the main olfactory system (MOS) and accessory olfactory system (AOS) are the two major sub-systems of the olfactory system (Beites *et al*., 2009). The MOS consists of the main olfactory epithelium that can detect odorants and send information to the main olfactory bulbs.

The AOS consists of the vomeronasal organ (VNO) that is responsible for a range of social and sexual behaviors as it can detect pheromones and kairomones and transmit the information to accessory olfactory bulbs (AOBs) (He *et al*., 2008; Papes *et al*., 2010). However, the VNO is comparatively less studied regarding stem cell heterogeneity and its regenerative ability (Katreddi and Forni, 2021).

Studies in the main olfactory epithelium (MOE) highlighted the presence of two distinct types of basal stem cell populations in the adult stages based on their morphology and position in the epithelium - Globose Basal Cells (GBCs) and Horizontal Basal Cells (HBCs) (Graziadei and Graziadei, 1979; Schwob and Jang, 2006; Schwob *et al*., 2017). The term GBCs includes a variety of mitotically active cell populations, including 1) multipotent Sox2/Pax6+ stem cells, 2) Ascl1+ transit amplifying neuronal progenitors, and 3) unipotent Neurog1/Neurod1+ immediate neuronal precursors that further divide and give rise to olfactory sensory neurons (Guo *et al*., 2010; Shaker *et al*., 2012).

HBCs are a second population of basal stem cells that lie juxtaposed to the basal lamina of the MOE. These cells constitute a stem cell reservoir that stays dormant in the intact adult olfactory epithelium. HBCs express p63 TF, specifically ΔNp63α isoform (Fletcher *et al*., 2011; Packard *et al*., 2011). p63 is a key factor in maintaining the stemness of epithelial stem cells (Li *et al*., 2023).

HBCs can be activated, proliferate, and differentiate to give rise to neuronal and non-neuronal cell types of the MOE within weeks of time after sensory epithelium lesions (Fletcher *et al*., 2017; Herrick *et al*., 2018). Interestingly, in rodents, the HBCs form at later stages of embryonic development compared to Sox2-positive stem cells seen as early as E10 (Beites *et al*., 2009; Packard *et al*., 2011). How the HBCs form is still a matter of investigation.

The VNO in mice is a bilaterally symmetrical tubular structure at the base of the nasal cavity responsible for detecting pheromones (Matsunami and Buck, 1997). It comprises a thick medial vomeronasal sensory epithelium (VSE) and a thin lateral non-sensory epithelium (NSE) with a lumen separating both epithelia (Katreddi and Forni, 2021; Takami, 2002).

Within the VSE of rodents, there are 2 main types of VSNs that are crucial in triggering a number of stereotypical behaviors in response to intra and inter specific pheromonal communication (Chamero *et al*., 2011; Chamero *et al*., 2012; Perez-Gomez *et al*., 2015; Trouillet *et al*., 2019).

The evolutionary, more conserved type of VSNs is formed by cells expressing the transcription factor Meis2. These neurons mostly express the Gαi2 subunit and receptors of the V1R family. Based on their spatial distribution, these neurons are often referred to as apical vomeronasal sensory neurons (VSNs) (Belluscio *et al*., 1999; Berghard and Buck, 1996; Buck and Axel, 1991; Del Punta *et al*., 2002; Enomoto *et al*., 2011; Trouillet *et al*., 2019).

The second type of VSNs, mostly abundant in rodents and marsupials, comprises neurons expressing the transcription factor Tfap2e/AP-2ε. These cells express the Gαo subunit, and receptors of the V2R family. These VSNs are often called basal VSNs (Chamero *et al*., 2011; Dulac and Axel, 1995; Herrada and Dulac, 1997; Matsunami and Buck, 1997; Ryba and Tirindelli, 1997).

Birth dating studies in the VNO indicate that postnatal neurogenesis mostly occurs at the marginal zones of the VSE. From there, the newly formed neurons migrate horizontally toward the central areas of the VNO (Brann and Firestein, 2010). However, pools of stem cells located at the level of the basal lamina at the center of the sensory epithelium have also been described and proposed to participate in cell renewal from their respective position. Neurons forming in the central regions have been described as undergoing a vertical migration (Brann and Firestein, 2010; Martinez-Marcos *et al*., 2005).

In terms of progenitor cells in the VNO, the role and turnover of the globose basal cell equivalents, which also express Sox2, Ascl1, Neurog1, and Neurod1 have been extensively characterized (Cau *et al*., 2002; Katreddi *et al*., 2022; Panaliappan *et al*., 2018). As these cells have nearly identical functions as the globose basal cells of the MOE, we will call them vomeronasal GBCs (vGBCs).

In a recent study, we found that Notch1-Dll4 signaling in vGBCs is important for establishing the diverging developmental trajectories of apical and basal VSNs (Katreddi and Forni, 2021; Katreddi *et al*., 2022).

A previous study reported the existence of Keratin-14 (Krt14) positive basal cells in the non-sensory epithelium (NSE) and marginal zone of VNO (Takami *et al*., 1995). However, what these cells are, has not been further explored.

Only a few studies have focused on the NSE of the VNO. This epithelium has been described both as stratified and as pseudostratified, depending on the animal species analyzed (Eltony and Elgayar, 2011; Garrosa *et al*., 1998).

In the current study, we identified p63+/Krt5+/Krt14+ basal cells in the non-sensory epithelium (NSE) of the VNO. These cells morphologically and molecularly differ from the columnar apical cells facing the lumen. The basal cells of the NSE are progenitor cells that give rise the columnar cells. Moreover, we found that the VNO, like the MOE (Fletcher *et al*., 2011) has p63+/Krt5+/Krt14+ HBCs along the basal lamina of the sensory epithelium (SE). These cells appear postnatally. The vomeronasal HBCs originate from progenitors along the basal lamina of the marginal zones, forming a continuum with the basal cells of the NSE.

## Results

### ΔNp63, Krt14, Krt5, and Sox2 identify various basal cell populations along the sensory and non-sensory epithelia of the VNO

The transcription factor p63, specifically the ΔNp63α isoform, is a common marker for basal progenitors of various epithelia, including the respiratory epithelia (Rock *et al*., 2009). ΔNp63α is one of the initial markers expressed by the horizontal basal cells (HBCs) in the MOE (Packard *et al*., 2011). We performed immunohistochemistry against ΔNp63 on postnatal VNOs. This staining highlighted p63 expression in the basal cells of the NSE and in spindled cells along the basal lamina of the vomeronasal sensory epithelium (VSE) (Fig. 1A).

**Figure 1:**
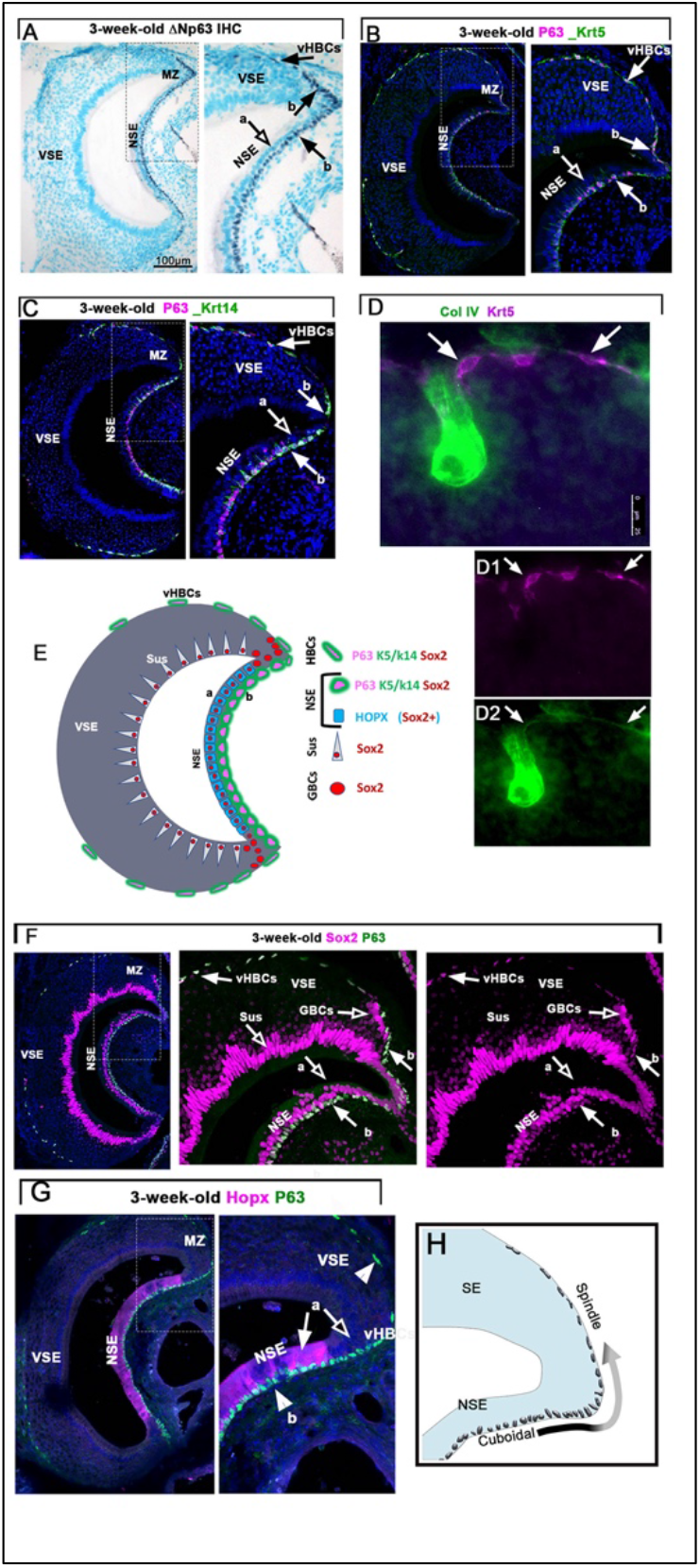
P63, Krt5 and Krt14 are expressed in the non-sensory epithelium of the VNO and by HBSc. **A).** P63 immunoreactivity (Ni-DAB dark gray; black arrow) in basal most ((b) Black arrow)) cells in the non-sensory epithelium (NSE). Apical cells of the NSE do not express p63 ((a) white arrow)). P63 is also expressed by vHBCs (black arrow) along the basal lamina of the Vomeronasal sensory epithelium (VSE). B) Immunofluorescence anti p63 (magenta) and Krt5 (green) shows the co-expression of both markers (arrow) in the basal most cells of NSE (White arrows). Empty arrows indicate apical cells in the NSE negative for the staining. K5 and p63 are co-expressed in vHBCs along the basal lamina of the sensory epithelium of the VNO. **C)** Immunofluorescence anti DNp63 (magenta) and Krt14 (green) shows the co-expression of both markers (arrow) in NSE and in the vHBCs. **D)** Immunofluorescence anti ColIV and Krt5 shows the vHBCs touching the basal lamina (arrow) of the SE. **E**) Cartoon summarizing protein expression in the different vomeronasal cells populations. **F)** p63(green) and Sox2 (magenta) shows Sox2 and p63 co-expression in the basal cells of the NSE and in the vHBCs (arrows). Sox2 expression was also found in the apical cells of the NSE and in the sustentacular cells (SUS) of the VSE. The globose basal cells (GBCs) are negative for p63. **G)** Hopx highlights apical columnar cells of the NSE in apical (a) portion of the NSE. The Hopx cells are negative for p63. Hopx expression abruptly ends as the apical cells arrive proximal to the marginal zone (black arrow). No Hopx was found in the p63+ basal cells of the NSE (a) nor in the vHBCs. **H**) Cartoon illustrating the change is shape of the basal cells from cuboid in the NSE to fusiform from along the SE.

Double immunofluorescence against ΔNp63/Krt5 and ΔNp63/Krt14 at 3 weeks after birth showed that the cells expressing p63 in both NSE and along the basal lamina of the VSE also expressed both keratins (Fig. 1B, C). The ΔNp63/Krt5 cells were found in the basal portions of the non-sensory epithelium and attached to the Collagen IV+ basal lamina of the sensory epithelium from the marginal to the central zones (Fig. 1D).

Sox2/ΔNp63 double immunofluorescence confirmed that the horizontal basal cells of the VNO (vHBCs) express a similar molecular profile (p63, Krt14, Krt5, and Sox2) to those reported for the HBCs of the MOE (Peterson *et al*., 2019).

### The globose basal cells are negative for p63

Sox2/p63 double immunostaining highlighted that the GBCs in the marginal zones, the sustentacular cells of the VNO, and the apical cells of the non-sensory epithelium were positive for Sox2 expression but negative for p63 immunoreactivity (See summary in Fig. 1E).

However, p63 and Sox2 double positive cells were found along the basal lamina of the non-sensory and sensory epithelium (Fig. 1F).

Moreover, the immunostaining against the homeobox gene (Hopx), which highlights cells of the NSE (Chang and Parrilla, 2016), (Fig.1G), labeled columnar cells in the most apical layers of the NSE but not p63+ the basal cells. Indicating that the NSE of rodents is a stratified epithelium.

Interestingly, the basal p63+ cells of the NSE and the vHBCs often formed a continuum but displayed different gross cell morphology. The basal cells of the NSE and the p63/Krt5+ proximal to the marginal zones have a cuboidal morphology with round nuclei while the vHBCs display a spindled shape with flattened/fusiform nuclei in close apposition to the basal lamina, as described for the HBCs of the MOE (Carter *et al*., 2004)(Fig. 1A,B,G,H).

### Characterization of the spatiotemporal developmental pattern of p63

Once we identified p63/Krt5 expression in the basal cells of the NSE of postnatal VNOs, we decided to look at p63 and Krt5 expression at earlier stages of development.

We analyzed p63 protein expression in VNO at E12.5. At this stage, p63 was broadly expressed by ectodermal epithelia, such as the developing skin, respiratory epithelia in the nose, and oral and tongue epithelia (Fig. 2A). Notably, parasagittal sections of the VNO showed p63 immunoreactivity in the rostral portion of the VNO juxtaposed to the putative RE with no reactivity in the rest of the VNO.

**Figure 2:**
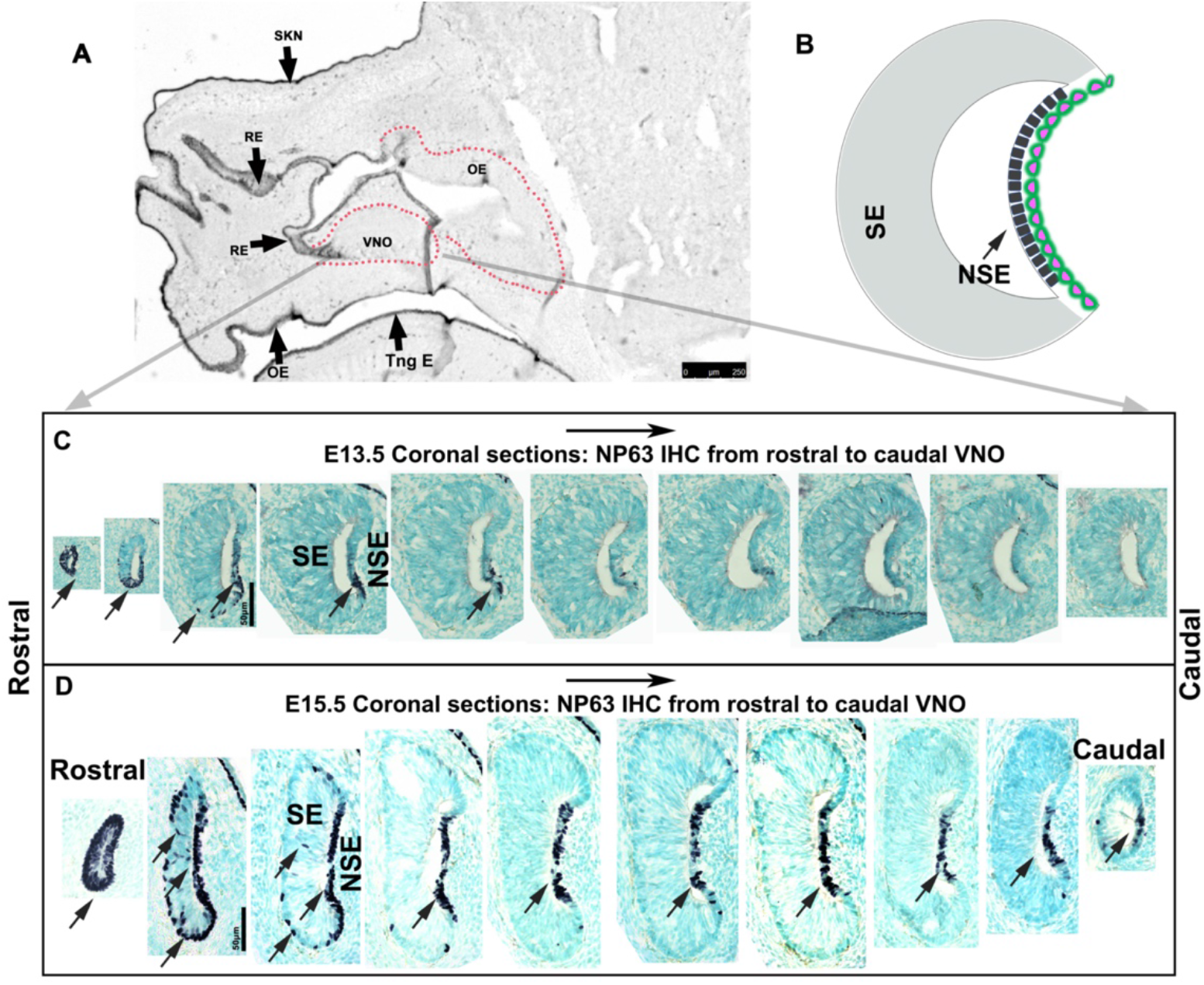
Dynamic expression of DNp63 from rostral to caudal VNO in early embryonic stages. **A**) E12.5 p63 expression is visible in the developing skin (Skn), respiratory epithelia (RE), oral epithelium (OE), and epithelium of the tongue (Tng E). **B**) Cartoon of coronal section of the VNO, sensory epithelium (SE), non-sensory epithelium (NSE). **C**) Expression of p63 (Arrows) from rostral to caudal sections of the VNO at E13.5 stage. p63 expression is mostly limited to cells in the rostral sections. **D**) p63 immunoreactivity from rostral to caudal sections at E15.5 stage. Arrows show the p63 expression in all sections of the VNO. Rostral sections show p63 cells in both sensory and non-sensory epithelia while caudal sections show p63 limited to the NSE.

Immunohistochemistry against p63 at E13.5 on coronal sections of the VNO showed some immunodetectable p63 in the rostral most sections but not in the caudal sections of the VNO (Fig. 2C). Intriguingly, by E15.5, p63 immunoreactivity could be found throughout the NSE from rostral to the caudal part of the VNO. Notably, at this stage, we found sparse p63 expression within the rostral most portions of the putative sensory epithelium (Fig. 2D).

### Krt5 expression follows p63 expression

Krt5 expression is controlled by p63 (Romano *et al*., 2009). Therefore, we exploited p63 and the downstream Krt5 expression to further follow p63-mediated gene expression dynamics over time.

p63 and Krt5 immunostainings at postnatal stages P0, P5, P20 and P30 showed immunoreactivity in the non-sensory epithelium and along the marginal zones of the VNO from birth. Quantification at different developmental stages showed that the number of p63 cells significantly increases in the first two weeks after birth. However, at birth very few of the p63 cells appeared to express Krt5. Krt5 immunoreactivity increased after P5 to reach an expression plateau around P15 in virtually all the p63+ cells (Fig. 3A-D, A’-D’).

**Fig. 3:**
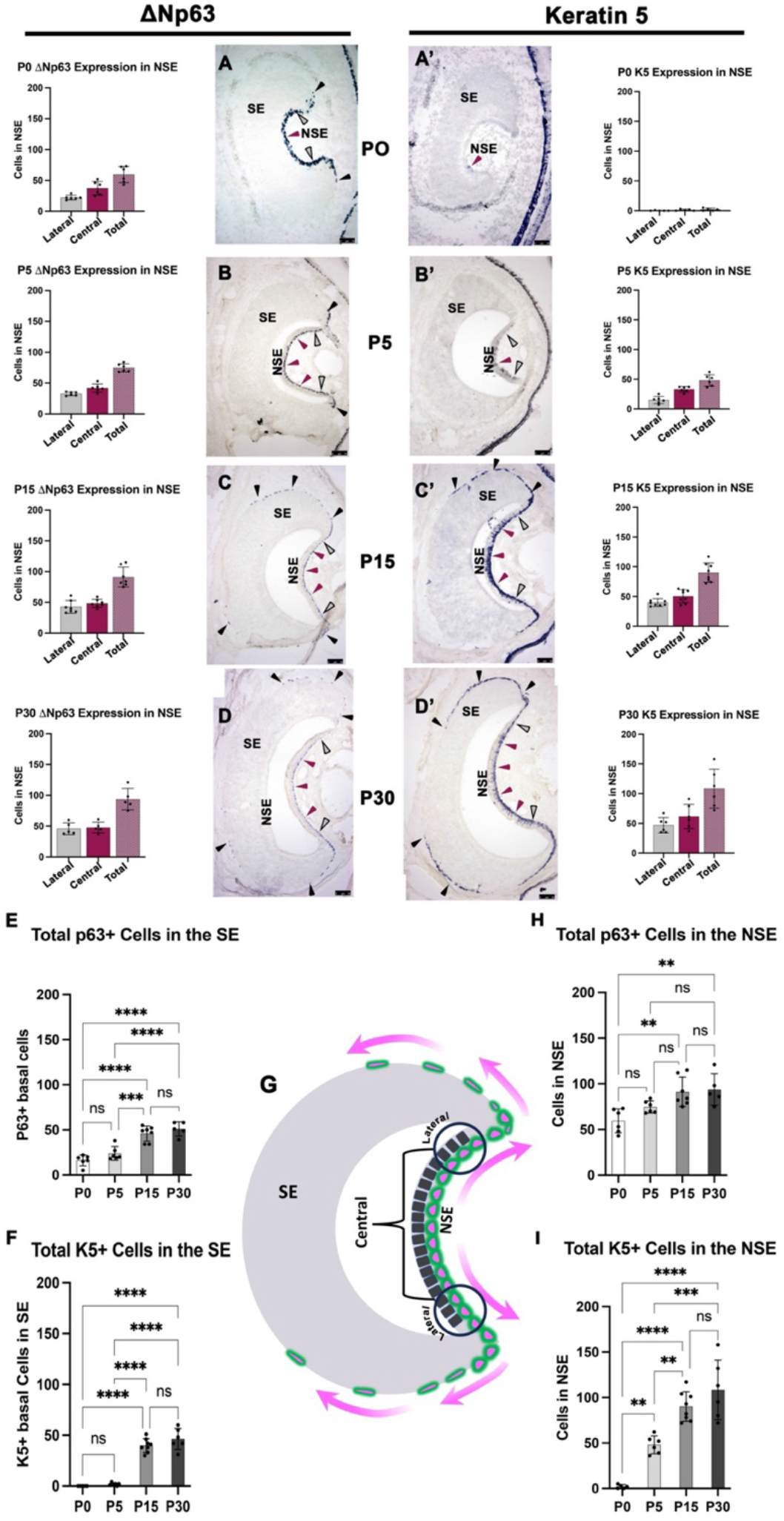
**A-D** Quantification of p63+ cells in the central and lateral regions of the NSE. Images from P0 to p30 show cells in the central (red arrows) and lateral regions of the NSE (white arrows). A progressive expansion of p63 immunoreactivity is visible along along the basal lamina of the SE (dark arrows).**A’-D’** Keratin 5 expression and quantifications in the central regions of the NSE (red arrow) and lateral regions of the NSE (gray arrows). **E**) Increase in p63+ cells in the SE from P0 to P30. **F**) Increase in Krt5+ cells in the SE form P0 to P30. **H, I**) Increase of p63+ and Krt5 cells in the NSE from P0 to P30. Values +/- STDEV; ANOVA followed by Tukey’s multiple comparisons test (*P <0.05). (**P<0.005). (***P<0.0005). **G**) Cartoon of coronal section of the VNO depicting the quantification regions and expansion of p63/Krt5 reactivity.

Following the expression of p63 and Krt5 in the central region and lateral regions of the NSE (Fig.3G), and along the SE, we found that Krt5 immunoreactivity progressively expands from the central areas of the NSE towards the lateral portions of the NSE, which are proximal to the marginal zones and then to the basal lamina of the SE (Fig. 3E, F, H, I). The dynamic changes in Krt5 expression over time suggested a potential expansion of the p63 progenitors from the NSE to the marginal zones and then to the central zones of the SE (Fig. 3G). As Krt5 immunoreactivity is delayed compared to p63, the dynamics of expression of Krt5 suggest that the progenitor cells giving rise to the vHBCs might be in the most lateral potions of the non-sensory epithelium proximal to the marginal zones.

### Sc-RNAseq profiling of the VNO identifies heterogeneous cell populations in the NSE

Analyzing merged scRNA-seq data from whole VNOs at P10, P21 and P60, we identified the cluster formed by the cells of the developing VSNs (Fig. 4A). This Y-shaped group of subclusters comprises the proliferative vGBCs (Ki67+, Sox2+, Ascl1+, Neurod1+) giving rise to the apical (Meis2; V1R) and basal (tfap2e; V2R) VSN dichotomy (Fig. 4B) VSNs (Katreddi *et al*., 2022). In a large neighboring group of clustered cells, we identified multiple subclusters of p63, Keratin5, and Krt14+ cells belonging to the putative NSE (Fig. 4A, isolated in 4C). Previous works suggested Icam1 and Itgb4 as HBCs preferentially expressed genes. However, these, as p63, Keratin5, and Krt14, were found to be broadly expressed across the cells of the putative NSE. We could not identify a definitive cluster of cells as the vHBCs.

**Figure 4:**
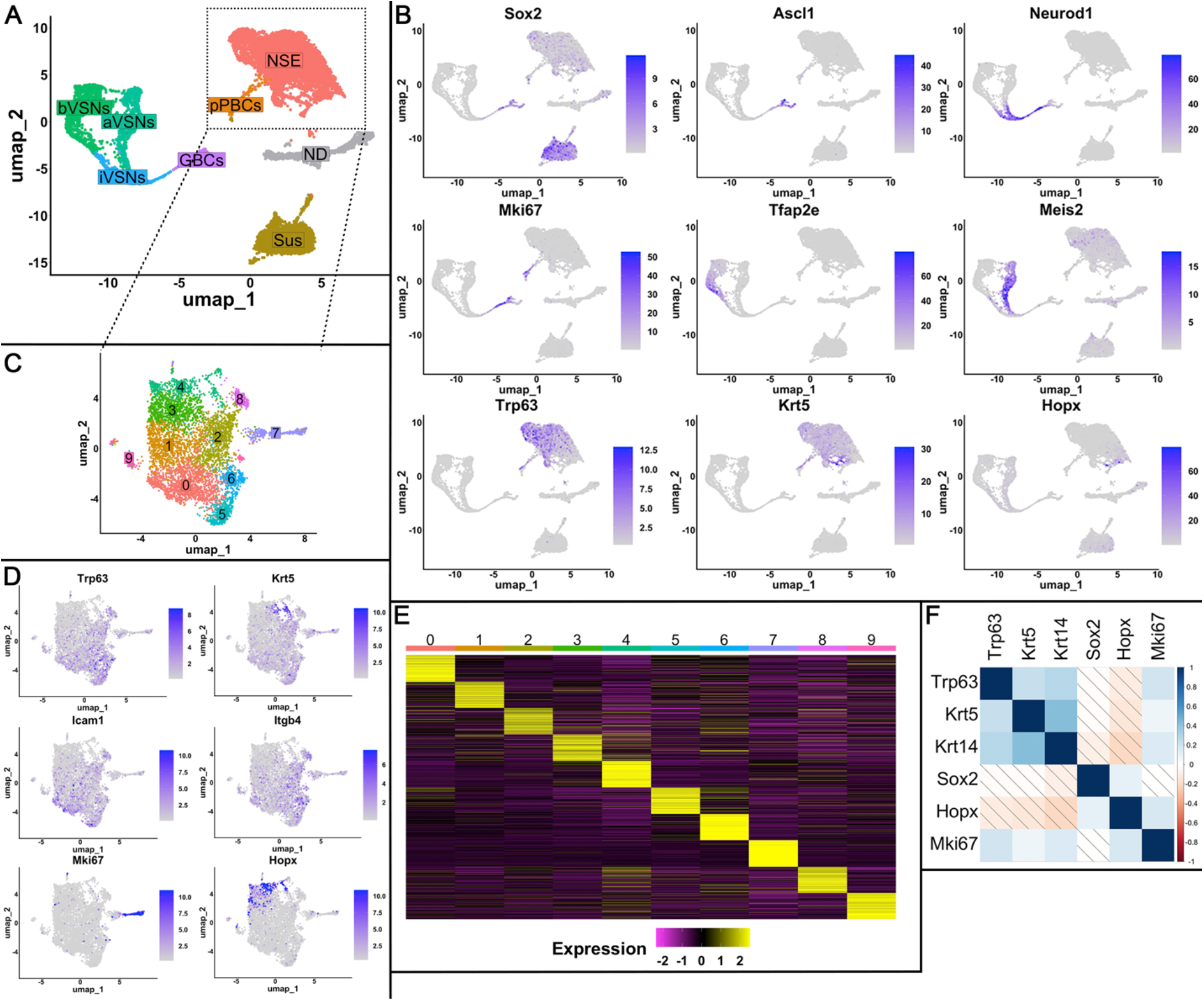
scRNA-seq identified NSE connection to VNO. **A)** UMAP dimensional reduction plot of Seurat object shows different cell types in the NSE and VNO. Each colored cluster of cells corresponds to an identified cell type that has a similar transcriptomic profile non-sensory epithelium (NSE), putative Proliferative Basal Cells (pPBCs); Sustentacular cells (Sus), Globose Basal cells (GBCs), Immature vomeronasal sensory neurons (iVSNs), apical VSNs (aVSNs), basal VSNs (bVSNs), non-determined (ND). **B)** Feature plots of Sox2, Ascl1, Neurod1, Mki67, Krt5, Trp63, Hopx and Cbr2 identify the VSN dichotomy and putative NSE. **C)** UMAP dimensional reduction plot shows the putative NSE without the cells of the VNO. **D)** Feature plots of Trp63, Krt5, Mki67, Icam1, Itgb4, and Hopx show the differential distribution of the Trp63/Krt5 and Hopx+ cells. **E)** Heatmap showing the top 100 differentially expressed genes in the 9 clusters shown in C. **F)** Correlation analysis of the genes expressed by the cells of the putative NSE, diagonal lines indicate no statistical correlation or negative correlation.

However, Hopx expression was found to be restricted to two subclusters of cells (Fig 4. C,D), mostly negative for the basal cell markers.

Performing a correlation analysis ((Seurat; corrplot); Fig.4F) using data from all the cells of the putative NSE we observed that the Hopx+ expression in the NSE appeared to correlate with decreased p63, Krt5, and Krt14 expression.

The proliferative marker Ki67 appeared enriched in subclusters of p63, Krt5, Krt14 (Fig. 4D,E). Gene expression profiling across clusters of the putative NSE indicates considerable genetic heterogeneity across cell clusters ((Fig.4 E); Supplementary Tab.1).

### Keratin5 lineage tracing

Performing immunolabeling against p63 and ki67 in P21 animals, we found, in line with what was described in other species (Elgayar *et al*., 2017), that, in postnatal animals, the proliferative cells of the NSE are, for the most part, basal progenitors (Fig 5A). These observations made us wonder if 1) the p63+ basal progenitors are the stem cells of the NSE and 2) if p63 basal cells could also be a source of other cell types in the VNO.

**Fig. 5.**
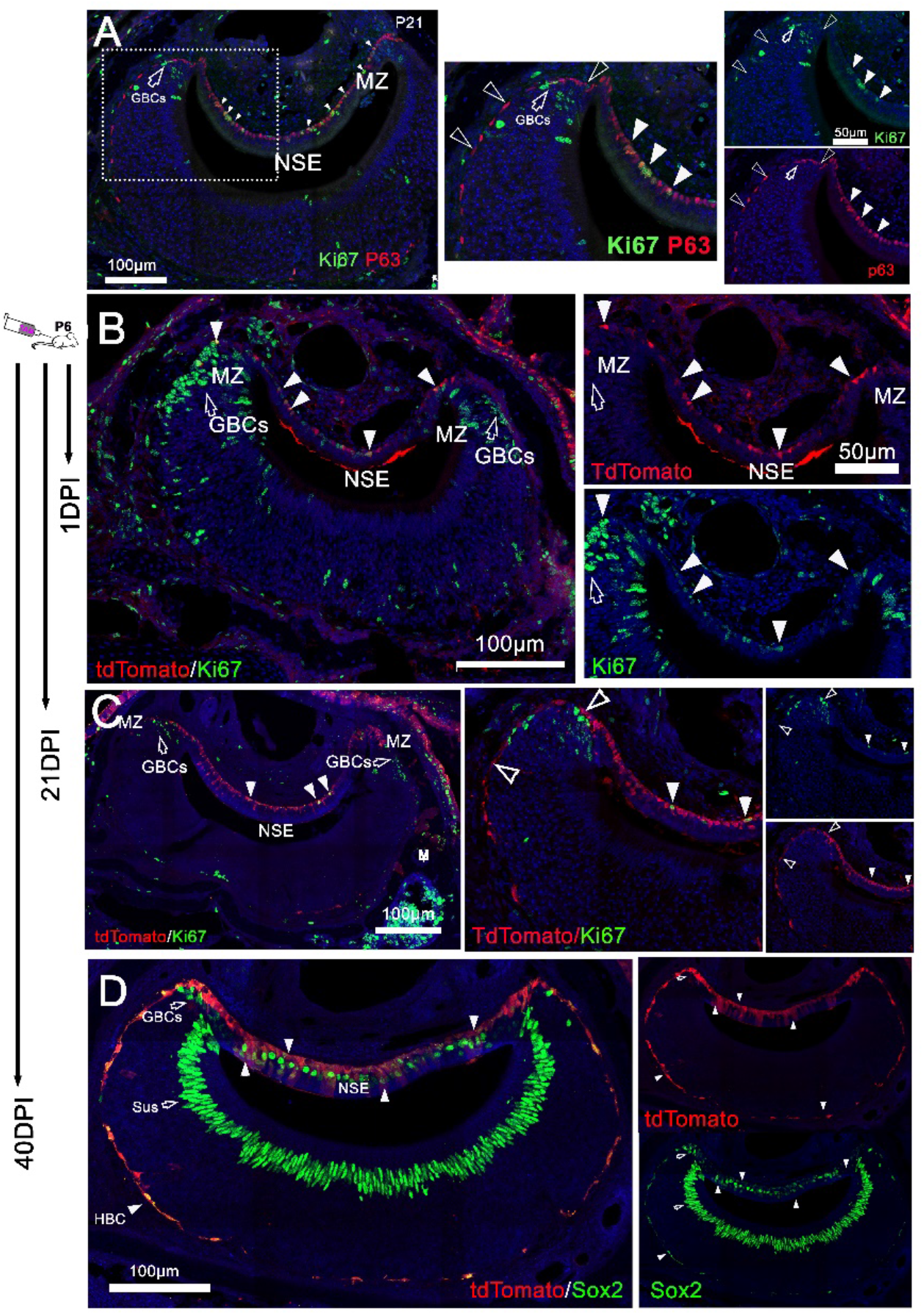
Proliferation of Krt5/p63 basal cells of the NSE. **A**) P21 p63 and ki67 show basal mitotic cells in the NSE positive for p63. The proliferative GBCs (empty arrows) in the Marginal zones (MZ) are negative for p63. **B**) VNO of Krt5CreERT2/R26tdTomato at P7(1DPI) shows Krt5Cre recombination (tdTomato) limited basal cells of the NSE. Ki67 (Green) is expressed by basal traced cells (Arrowheads). The proliferative GBCs (empty arrows) in the Marginal zones (MZ) are negative for Krt5 tracing. The traced cells along the SE (empty arrowheads) are not proliferative. **C**) P27 (21DPI), Ki67 is limited to the basal traced cells in NSE. The GBCs (empty arrows) in the MZ are negative for Krt5 tracing. D) At P46 (40DPI) Krt5 tracing is visible in basal and apical cells of the NSE, and in the HBCs along the basal lamina of the SE. However, Sox2+ GBCs and Sustentacular cells (Sus) appear negative for the tracing.

Krt5 is downstream p63 (Romano *et al*., 2009) therefore, its expression is delayed compared p63.

In the VNO Krt5 is restricted to the basal cells of the NSE and to putative vHBCs. Based on this, we decided to follow the lineage of p63/Krt5 cells in postnatal animals using a tamoxifen (TAM) inducible Krt5CreERT2 knock-in mouse line (Van Keymeulen *et al*., 2011).

Krt5CreERT2^+/-^/R26^tdTom+/-^ pups were injected with TAM at P6 and then euthanized for histological analysis at 1-day post injection (DPI), 14 D.P.I, 21 D.P.I and at 40 D.P.I. At 24h after injection, all the recombined (tdTomato+) cells were also positive for Krt5 (Fig 6 A3). At 1DPI, recombination was restricted to the basal cells of NSE and in sparse cells along the basal lamina of the marginal zones (Fig 5B).

**Fig. 6:**
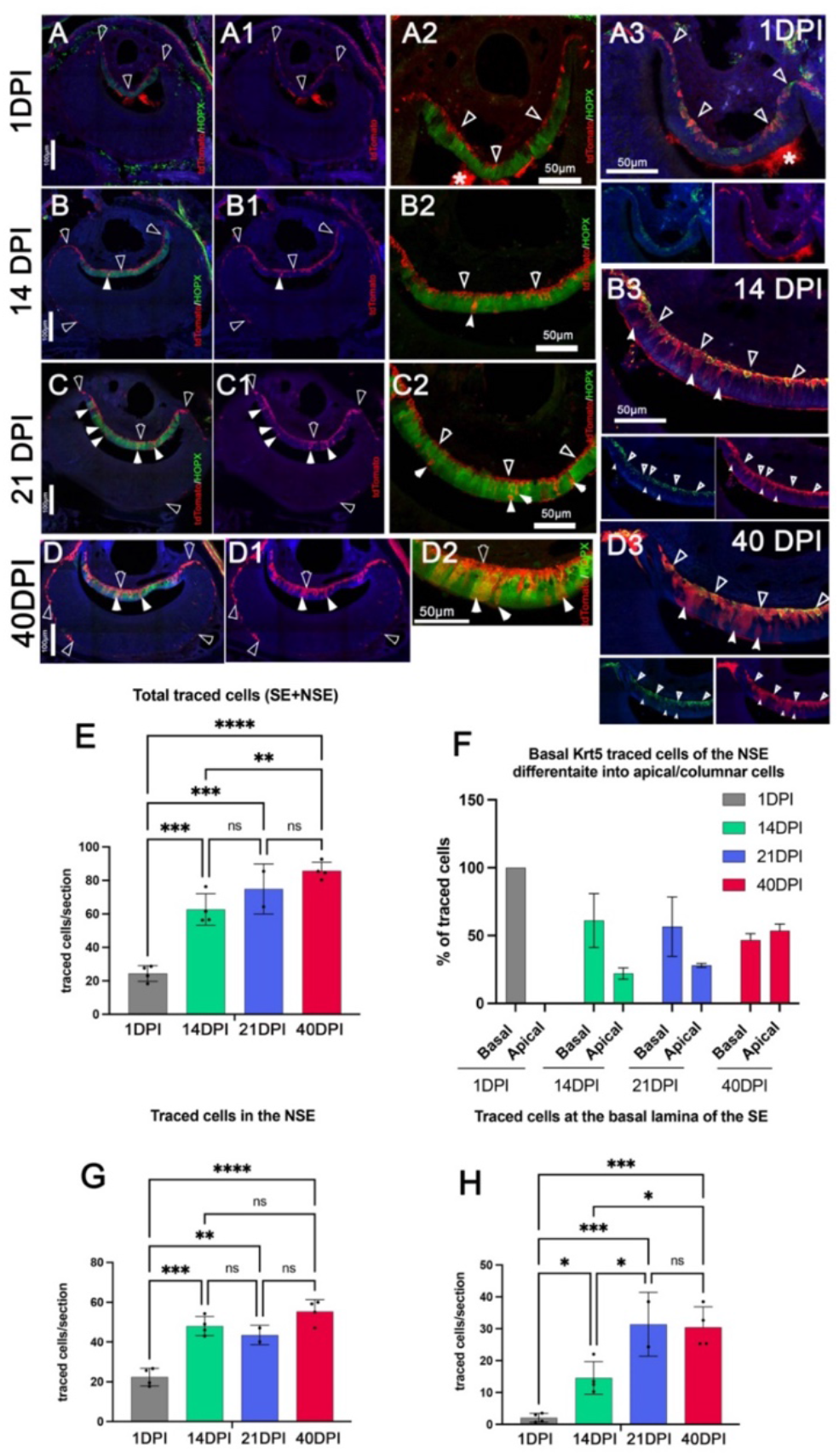
The basal cells of the NSE give rise to the columnar cells. Krt5 tracing at 1DPI (**A-D2**), 14DPI,21DPI and 40DPI. At 1 DPI the basal traced cells (empty arrowheads) are negative for Hopx. B-D2) At 14 DPI, traced cells differentiate into Hopx+ cells (White arrowheads) **A3-D3**) Traced cells in the apical regions (Empty arrowheads) of the NSE are positive for Krt5, while traced cells in the acpical region appear negative. **D3)** At 40DPI virtually all the basal Krt5+ cells derive from the traced progenitors. **E)** Increase in the number of traced cells between 1 and 40DPI. **F)** Changes in the distribution of traced cells between basal and apical regions of the NSE. **G)** Increase in traced cell numbers in the NSE. **H)** Increase in cell numbers along the basal lamina of the SE. Values +/- SD, ANOVA followed by posthoc Tukey’s multiple comparisons test (*P <0.05). (**P<0.005). (***P<0.0005)

Ki67 paired with tdTomato immunostaining confirmed that the basal cells of the NSE are proliferative progenitors.

Quantifications at 1DPI indicated that at P7 ∼20% of the traced cells in the NSE were proliferative (ki67+). However, along the basal lamina of the marginal zones only 4% of the traced cells resulted Ki67 positive. Notably, the globose basal cells in the marginal zones were negative for the tracing.

At 14DPI 13% of the traced cells in the NSE still resulted to be proliferative, however at this stage none of the traced cells along the marginal zones or along the basal lamina of the SE resulted to be proliferative.

Analysis at 40 DPI (Fig 5B-D) revealed that the Krt5 tracing was limited to the NSE and to the HBCs along the basal lamina of the SE. The Sox2+ globose basal cells were negative for the tracing.

### Basal cells differentiation in the NSE

Immunostaining against Hopx at 1DPI showed that the traced basal cells of the NSE and the Hopx-positive apical columnar cells of the NSE formed two distinct cell layers (Fig. 6A). However, between 21DPI and 40 DPI a larger proportion of the traced cells was recognizable as columnar cells in the apical regions of the NSE (Fig. 6A3-D3). By 40DPI around 50% of the cell positive for tracing could be found as differentiated columnar cells in the apical layer of the NSE.

At 40 DPI, most of the Krt5+ basal cells of the NSE still appear to be traced. These data suggest that the basal Krt5+ cells undergo both self-renewing symmetric divisions and non-symmetric divisions, differentiating into to columnar apical Hopx+ cells. Moreover, the increase in number of Krt5 traced cells, from 1DPI to 40 DPI, along the basal lamina of the sensory epithelium suggests that the vHBCs originate from the basal Krt5+ progenitors at the border between the NSE and SE.

### Some of the things that did not go as expected

In our attempts to follow the lineage of the p63/Krt5 cells in the VNO we also tested the ΔNp63Cre mouse line (Pignon *et al*., 2013). These mice have been previously reported to give faithful recombination in ΔNp63+ cells but with low and chimeric efficiency (Pignon *et al*., 2013). Performing constitutive ΔNp63 tracing with this mouse line we observed highly chimeric recombination in the VNO with only sparse cells recombining the NSE and occasional tracing of the HBCs. Surprisingly, constitutively active ΔNp63Cre recombined in neurons and sustentacular cells in the MOE (not shown) and in the VNO (Supplementary figure 1). The tracing obtained with this mouse does not recapitulate p63 expression in the VNO.

To further investigate whether the vHBCs of the VNO have similar regenerative ability to that of the MOE, we tried to use methimazole intraperitoneal injection, a very well-established injury model in the MOE (Lin *et al*., 2017). We injected methimazole at 75mg/Kg bd wt and perfused at 3 days post-injection. Staining against cleaved caspase3 showed cell death only in the MOE but not VNO (Supplementary Figure 2), making this approach unsuitable for testing the regenerative ability of the vHBCs.

## Discussion

P63 is one of the two mammalian paralogues of p53 family of transcription factors that can exist in 6 different isoforms, two different N terminal, and 3 different C terminal isoforms (Ou *et al*., 2007; Yang *et al*., 1998). Specifically, the N-terminal truncated (ΔN) p63 isoform has been reported in the basal cells of different stratified epithelia (Rock *et al*., 2009) and has been shown to have a role both during development and homeostasis (Packard *et al*., 2011; Pignon *et al*., 2013; Yang *et al*., 1999).

In the MOE, which is a pseudostratified neuroepithelium ΔNp63 has been identified in the HBCs (Schnittke *et al*., 2015). These are a quiescent stem cell population with a role in post-injury regeneration (Fletcher *et al*., 2011; Packard *et al*., 2011) and in the cells of the respiratory epithelium.

Even though the VNO shares multiple molecular and morphological features with the MOE, it has been less characterized regarding stem cell populations. Several studies have focused on the developing SE while the NSE has been largely overlooked.

In this paper, we described 1) the existence of HBC in the VNO and 2) we followed the lineage of basal p63+/Krt5+ progenitors of the NSE.

Analysis of embryonic samples showed that p63 is first expressed in the developing VNO in the cells of the putative NSE. Krt5 expression, which follows p63 expression, only becomes detectable as a protein at early postnatal stages.

Analyzing p63 expression over time, we observed that the VNO has, like the MOE, fusiform HBCs underneath the basal lamina of the SE. The vomeronasal HBCs are positive for the HBC markers (Carter *et al*., 2004; Packard *et al*., 2011) such as p63, Sox2, Krt5, Krt14. Notably, these are also markers shared with the basal cells of the nasal respiratory epithelia (Davis and Wypych, 2021).

After birth, we observed a progressive increase in number of HBCs around the sensory epithelium that appear to migrate to the central zones of the VNO starting from basal cells lining proximal to the marginal zone.

Krt5 immunolabeling and genetic tracing confirmed that the HBCs originate from proliferative Krt5+ basal progenitors at the border between the NSE and the basal lamina of the marginal zones.

P63 is a key factor in maintaining the stemness of epithelial stem cells (Davis and Wypych, 2021; Li *et al*., 2023). The proximity between the vomeronasal HBCs and the basal cells of NSE suggests a potential direct lineage between these cell types. This idea is partially supported by our genetic lineage experiments. In fact, In Krt5CreER traced animals at P7 (1DPI), we could detect obvious recombination in the NSE but only sparse cells along the basal lamina of the marginal zones. However, analysis at 14, 21 and 40 DPI showed an overall increase in the number of traced cells in the NSE and traced HBCs throughout the basal regions of the SE. Once along the basal lamina of the SE the vHBC do not appear to be proliferative. These data indicate that the HBCs originate from proliferative Krt5+ progenitors at the marginal zones proximal to the NSE.

Analyzing scRNA seq data, we could not discriminate between basal cells of the NSE and HBCs because virtually all the previously identified HBCs markers (Carter *et al*., 2004) appear to be shared with cells of the NSE.

It is tempting to speculate that p63 basal cells forming in the NSE are stem cell that retain a sufficient level of potency to acquire differential regenerative potential based on their localization along the basal membrane of either neuronal or non-neuronal epithelia. Similar developmental studies focusing on the lineage and genesis of the HBCs (Fletcher *et al*., 2017) of the main olfactory epithelium could confirm this. Notably, the HBCs of the MOE are already formed at perinatal stages. Our experimental paradigms at postnatal stages did not allow us to follow their development as part of this work Future investigations focusing on identifying the expression of the most differentially expressed genes across clusters of cells that we identified via Sc-RNAseq (Fig.4) could reveal new cell populations or interesting transitional states of the cells of the NSE of the VNO.

Conditional Krt5 tracing of the basal cells of the NSE showed that the basal Krt5 + cells give rise to the columnar apical cells of the NSE. These non-surprising results indicate that the NSE of the VNO of rodents is, like respiratory epithelia (Rock *et al*., 2009), a stratified epithelium where basal progenitors (Sox2, Krt5, p63+) can give rise to columnar apical cells positive for Hopx.

One aspect we were not able to address in this study is whether the HBCs of the VNO have regenerative abilities in response to lesions as the ones of the MOE.

Previous studies have shown that tissue lesions or conditional LOF of p63 in HBCs of the MOE induce activation of the HBCs and tissue regeneration. However, in our hands, all Krt5CreERT/p63^Flox/Flox^ animals invariably died 1 short after TAM treatment, making it impossible for us to test the regenerative ability of the vHBCs (data not shown). Moreover, we also failed in our attempts to induce tissue damage in the VNO using Methimazole (Supplementary Figure 2). In fact, after Methimazole treatments, we observed considerable OE damage but no cell death in the VNO (Supplementary figure 2).

The lack of cell death in the VNO of methimazole-injected animals could result from quantitative and spatial differences in the expression of cytochrome P450 enzymes between the MOE and VNO. These enzymes are essential for the formation of toxic intermediate metabolites of methimazole (Chen *et al*., 1992; Gu *et al*., 1999). It is previously shown that cytochrome P450 enzymes are lowly expressed in the VNO (Gu *et al*., 1999) and at least in the rat VNO, cytochrome P450 enzymes were detected in intraepithelial duct cells and in the mucus layers above the sensory and non-sensory epithelia unlike in the sustentacular cells of the MOE (Chen *et al*., 1992).

If the newly identified HBCs of the VNO have regenerative potential, still needs to be addressed in future studies. Bulbectomy (Iwai *et al*., 2008) could be the right way to test this in the future.

## Author contributions

PEF conceptualized and designed the experiments and illustrations. NML and RRK designed, performed the experiments and analyzed the data, ND and NM performed and analyzed P21 RNA-seq experiments. NML, RRK, and PEF wrote the manuscript.

## Acknowledgments

We thank Carmela Polizzi for helping with sectioning and staining and Rico Amato for helping upload the RNA-seq data in GEO.

## Materials And Methods

### Mouse lines

The mouse lines Tg (KRT5-cre/ERT2)2Ipc/JeldJ Strain #:018394 (Indra *et al*., 1999); Krt5CreERT2 knock-in (B6N.129S6(Cg)-Krt5tm1.1(cre/ERT2)Blh/J Strain No: 029155 (Van Keymeulen et al., 2011), ΔNp63Cre knock-in (B6.129S-Trp63tm1.1(cre)Ssig/ Strain No: 024564) (Pignon *et al*., 2013), R26tdTom (B6.Cg-Gt(ROSA)26Sortm9(CAG-tdTomato) Hze/J, Strain No: 007909) were purchased from Jackson Lab. Mice of either sex were used for immunohistochemistry and immunofluorescence experiments. All experiments involving mice were approved by the University at Albany Institutional Animal Care and Use Committee (IACUC).

### Tamoxifen Treatment

Tamoxifen (Sigma–Aldrich), CAS # 10540−29−1, was dissolved in Corn Oil at 20ug/μl concentration. For all tamoxifen-inducible experiments, we injected tamoxifen once intraperitoneally at a dose of 75mg/Kg body weight and perfused at indicated postnatal days.

### Tissue Preparation

Tissue collected were perfused with PBS followed by 3.7% formaldehyde in PBS. Noses were immersion fixed in 3.7% formaldehyde in PBS at 4°C overnight. Noses were then decalcified in a 500 mM EDTA solution in PBS for a number of days proportional to their age. All samples were cryoprotected in 30% sucrose in PBS overnight at 4°C then embedded in Tissue-Tek O.C.T. Compound (Sakura Finetek USA, Inc., Torrance CA) using dry ice, and stored at -80°C. Tissue was cryosectioned using a CM3050S Leica cryostat at 18μm for VNOs and collected on VWR Superfrost Plus Micro Slides (Radnor, PA) for immunostaining. All slides were stored at -80°C until ready for staining.

### Immunofluorescence

Citrate buffer (pH 6.0) antigen retrieval was performed (Lin *et al*., 2018), for all the antibodies indicated with asterisks (*). Primary antibodies and concentrations used in this study were, *Rabbit anti-Ki67 (1:1000, D3B5, Cell signaling Tech), *Mouse anti-Ki67 (1:500, 9449, Cell signaling Tech), *Mouse anti-DsRd (1:500, TA180084, Origene), *Rabbit anti-DsRd (1:500, 600-401-379, Rockland), *Rabbit anti-Neurogenin-1 (1:400, 272926, abcam), *Goat anti-Sox2 (1:800, AF2018, R&D Systems), *Chicken anti-Keratin5 (1:1500, 905901, Biolegend), Chicken anti-Keratin14 (1:1000, 906001, Biolegend), *Rabbit anti-ϕλNp63α (1:500, 619002, Biolegend), *Rabbit anti-Hopx (1:200, 11419-1-AP, proteintech), Goat anti-CollagenIV (1:800, AB769, Millipore), *Rabbit anti-caspase 3 active cleaved (1:1000, AB3623, Millipore).

For chromogen-based reactions, tissue was stained as previously described (Lin *et al*., 2018). Staining was visualized with the Vectastain ABC Kit (Vector, Burlingame, CA) using diaminobenzidine (DAB); sections were counterstained with methyl green.

Species-appropriate secondary antibodies conjugated with either Alexa Fluor 488, Alexa Fluor 594, Alexa Fluor 568, Alexa Fluor 680 plus were used for immunofluorescence detection (Molecular Probes and Jackson ImmunoResearch Laboratories, Inc., Westgrove, PA). Sections were counterstained with 4’,6’-diamidino-2-phenylindole (DAPI) (1:3000; Sigma-Aldrich) and coverslips were mounted with FluoroGel (Electron Microscopy Services, Hatfield, PA). Confocal microscopy pictures were taken on a Zeiss LSM 710 microscope. Epifluorescence pictures were taken on a Leica DM4000 B LED fluorescence microscope equipped with a Leica DFC310 FX camera. Images were further analyzed using FIJI/ImageJ software.

### scRNA-seq quality control and cell clustering

P10, P21, and P60 scRNA-Seq data were integrated and processed to get a representation of different stages of development together. Quality control (QC), and clustering and downstream analysis was performed using Seurat (3.2.3) package in R. Basic filtering was carried out in which all genes that were expressed in ≥3 cells and all cells with at least 200 detected genes were included. QC was based on number of genes and percentage of mitochondrial genes – all cells that expressed >6665 genes, < 750 genes, <50,000 ncount_RNA and >10% mitochondrial genes were not included in the analysis. We used sctransform package to normalize and scale the data. Top 3000 highly variable genes across the population were selected to perform principal component analysis (PCA) and the first 40 principal components were used for cell clustering, which was then visualized using UMAP. Stem cells, neuronal progenitors, precursors and immature neuronal cell types were identified based on the expression of known genes. We removed one cluster, as it expressed genes of more than one cell type. Overall, 14,360 cells were included for the clustering and analysis.

### Experimental design, quantification, and statistical analyses of microscopy data

All data were collected from mice kept under similar housing conditions in transparent cages on a normal 12 hr. light/dark cycle. Tissue collected from either males or females in the same genotype/treatment group were analyzed together unless otherwise stated; ages analyzed are indicated in text and figures. Measurements of VNE and cell counts were performed on confocal images of coronal serial sections immunostained for the indicated targets. Measurements and cell counts were done using ImageJ. Quantification of postnatal chromogen-based reactions were completed by measuring the limit of the NSE to the center point of the NSE for both sides. The limit of the NSE was determined by placing a line tangential to the apical lamina of the NSE. A circle was then drawn equivalent in diameter to one quarter of the total NSE length. The diameter of the proportional circle was aligned along the NSE from the NSE limit inwards, this was repeated for both sides of the VNO. Immunoreactive cells within the bounds of the circles were considered lateral, while all remaining NSE were considered central. The data are presented as mean ± SEM unless otherwise specified. Prism 10.0.3 was used for statistical analyses, including calculation of mean values, and SEM. Two-tailed, unpaired t-test were used for all statistical analyses, and calculated p-values <0.05 were considered statistically significant. Sample sizes and p-values are indicated as single points in each graph and/or in figure legends.

### Data Availability

The scRNA-seq data for (P10) mice are available in GEO under accession number GSE192746. The scRNA-seq data for (P21) mice are available in GEO under accession number GSE247872. The scRNA-seq data for (P60) mice are available in GEO under accession number GSE190330.

---

**Supplementary figure 1.**
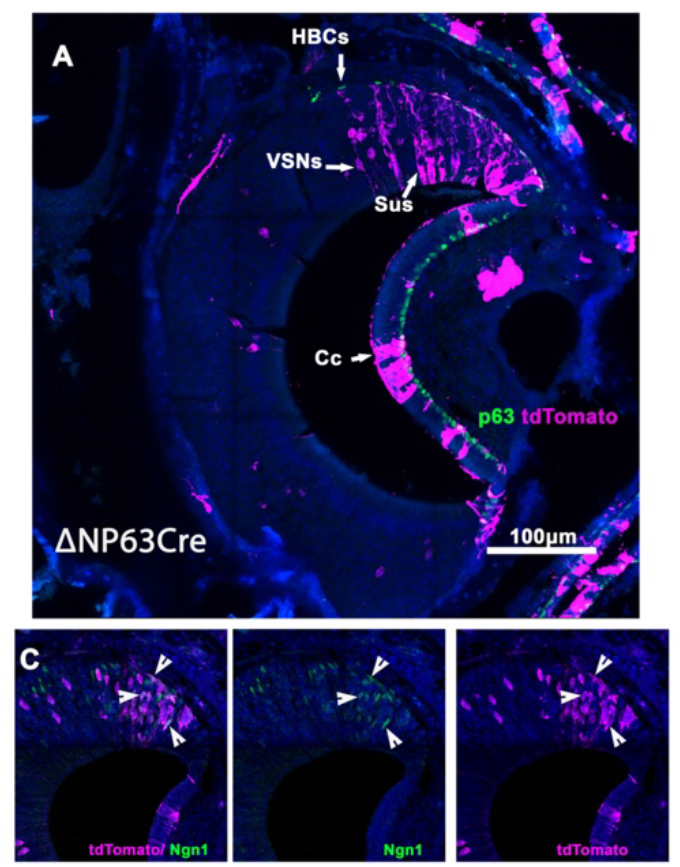
P30. **A**) Constitute p63 Cre tracing using DNp63Cre shows recombination only in sparse p63+ basal cells and columnar cells (Cc) in the NSE. However, recombination was observed in some VSNs and sustentacular cells (Sus). **C**) Neuronal progenitors positive for neurogenin1 and tracing.

**Supplementary figure 2.**
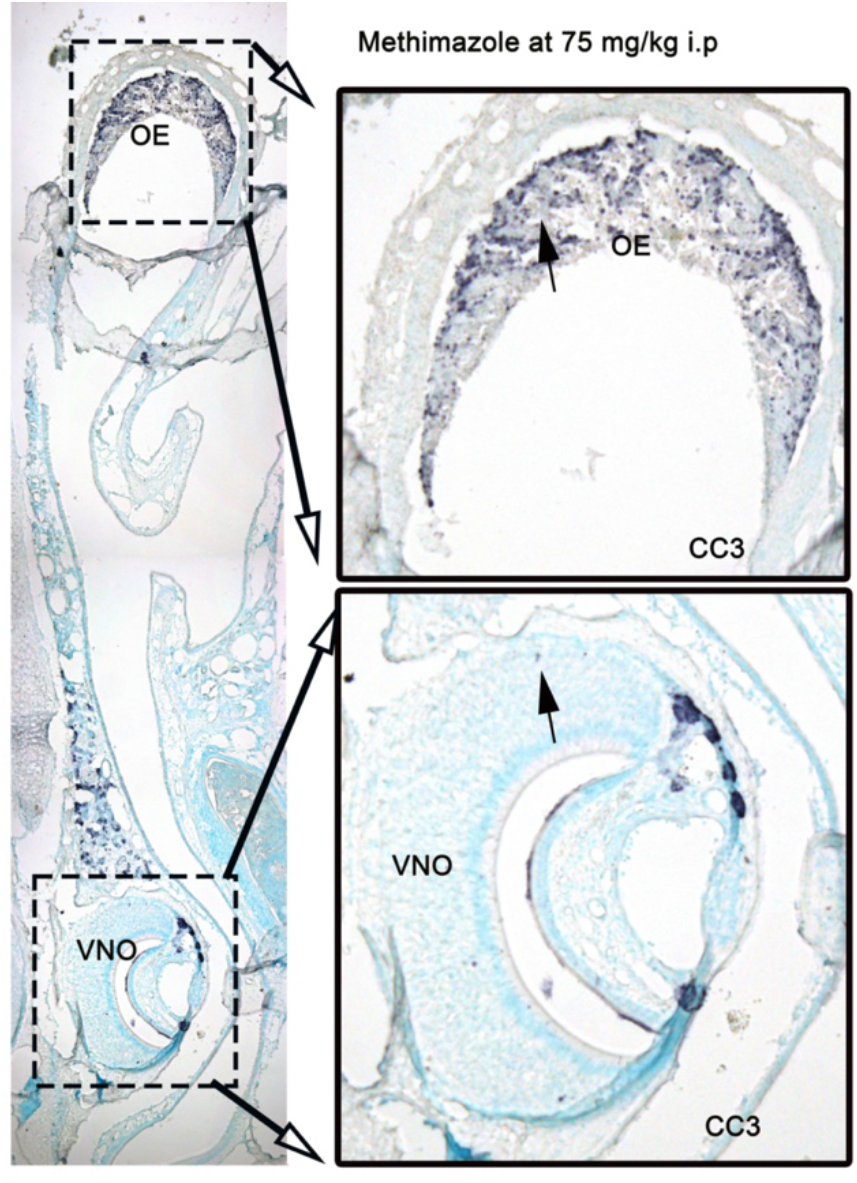
Adult WT animals, 3 days after IP injection of 75mg/Kg methimazole. Cleaved caspase staining (CC3) shows massive cell death in the olfactory epithelium (OE) but no cell death or visible tissue damage in the VNO.

**Supplemetary Table1.**
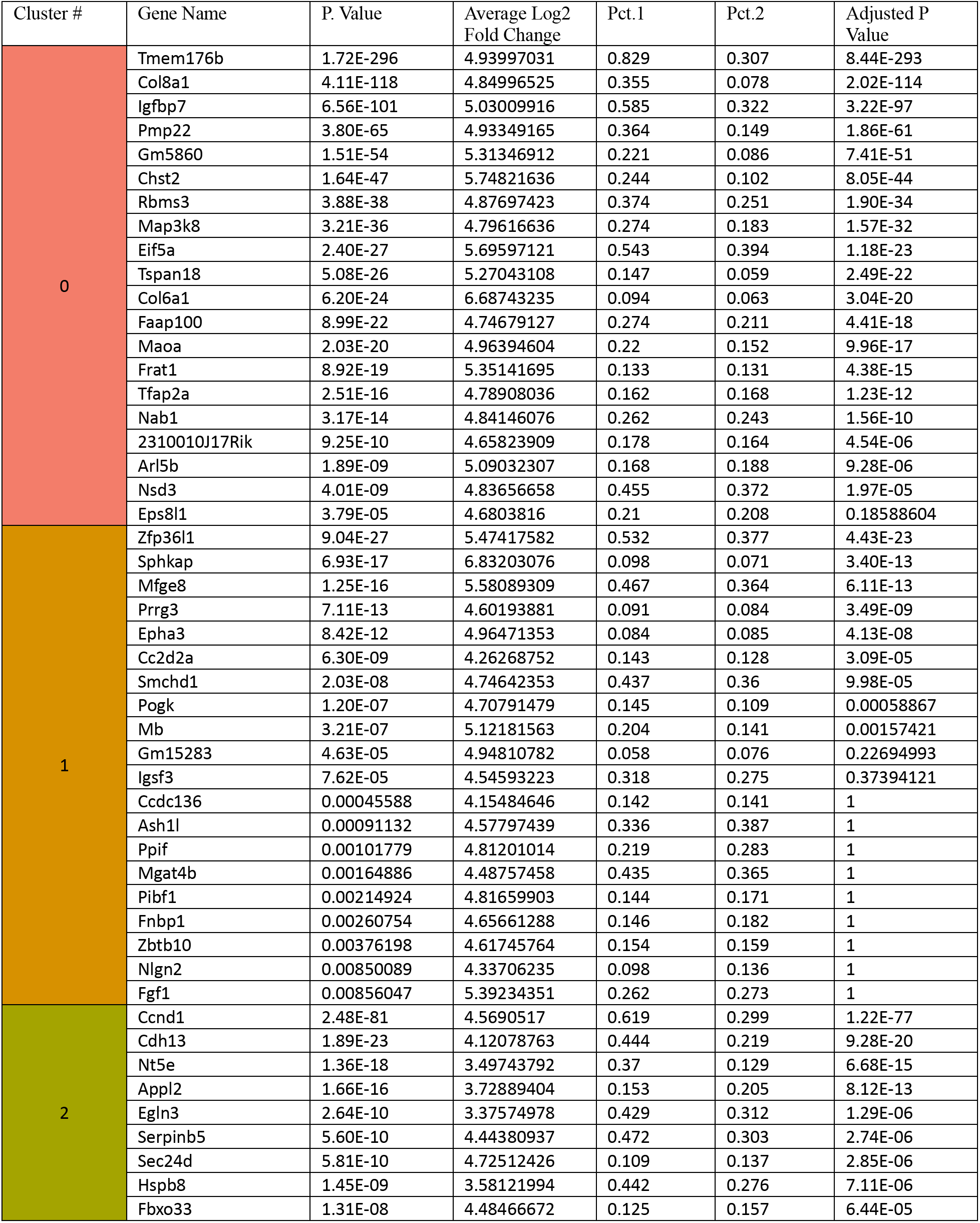
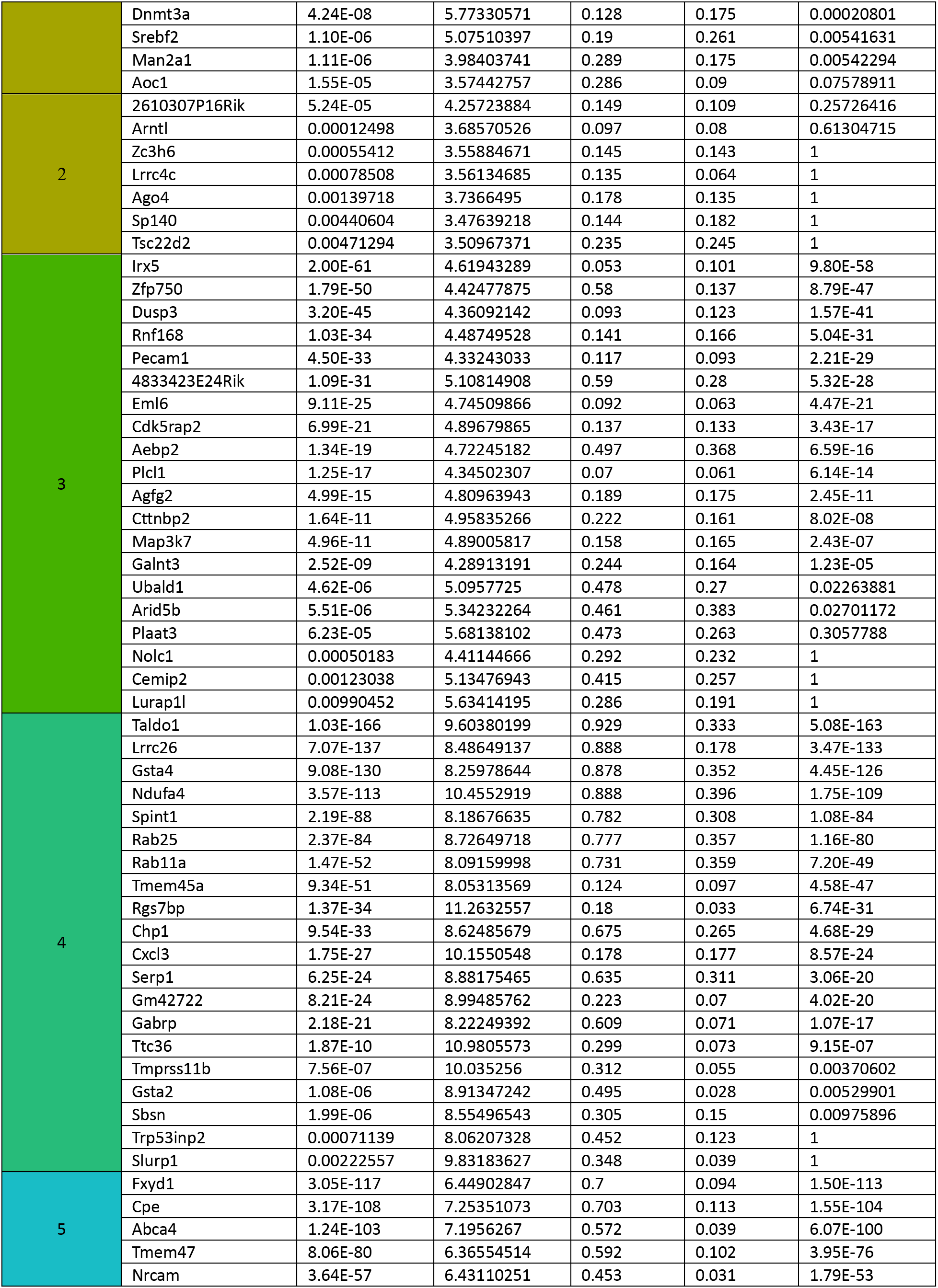
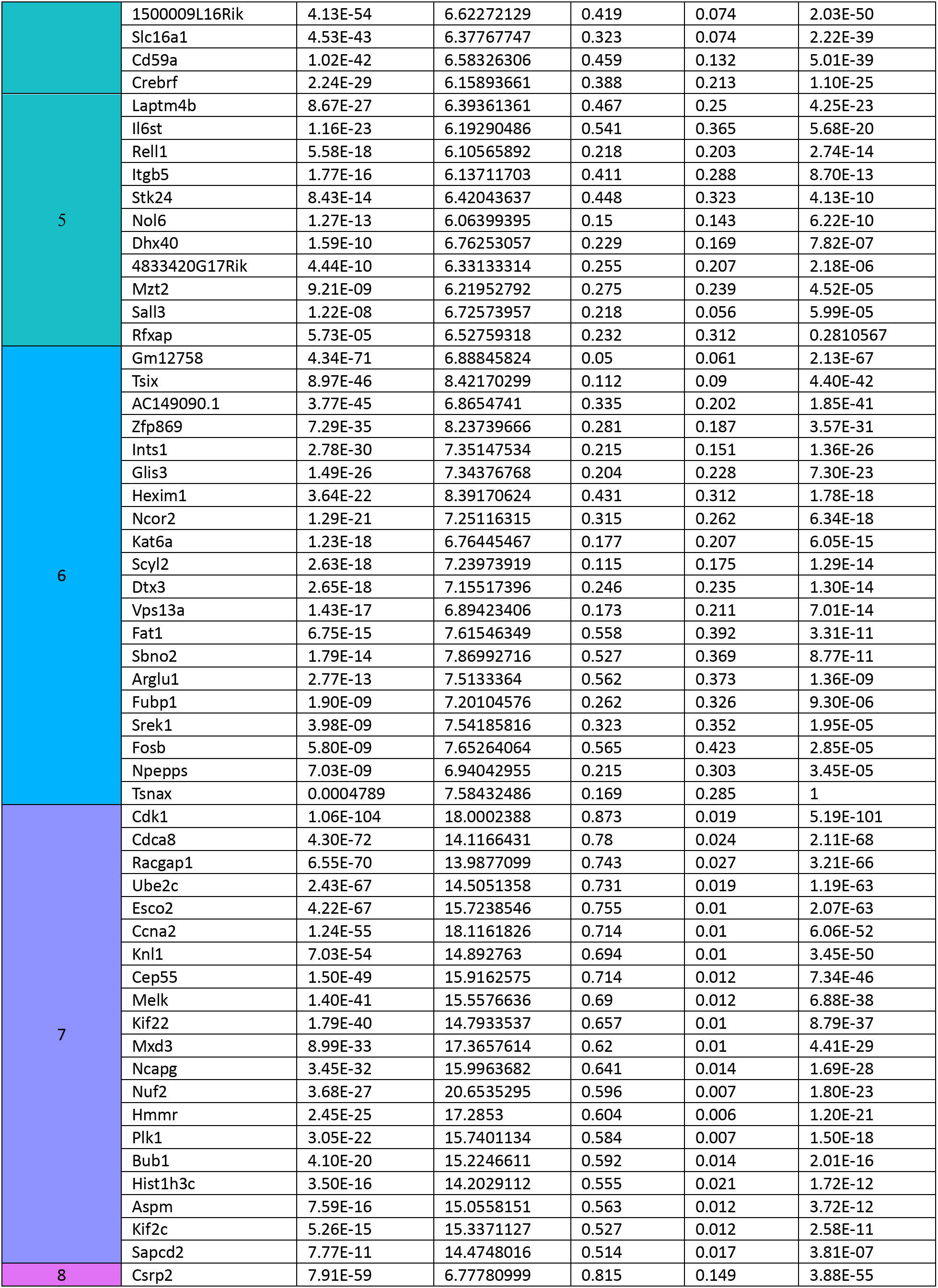
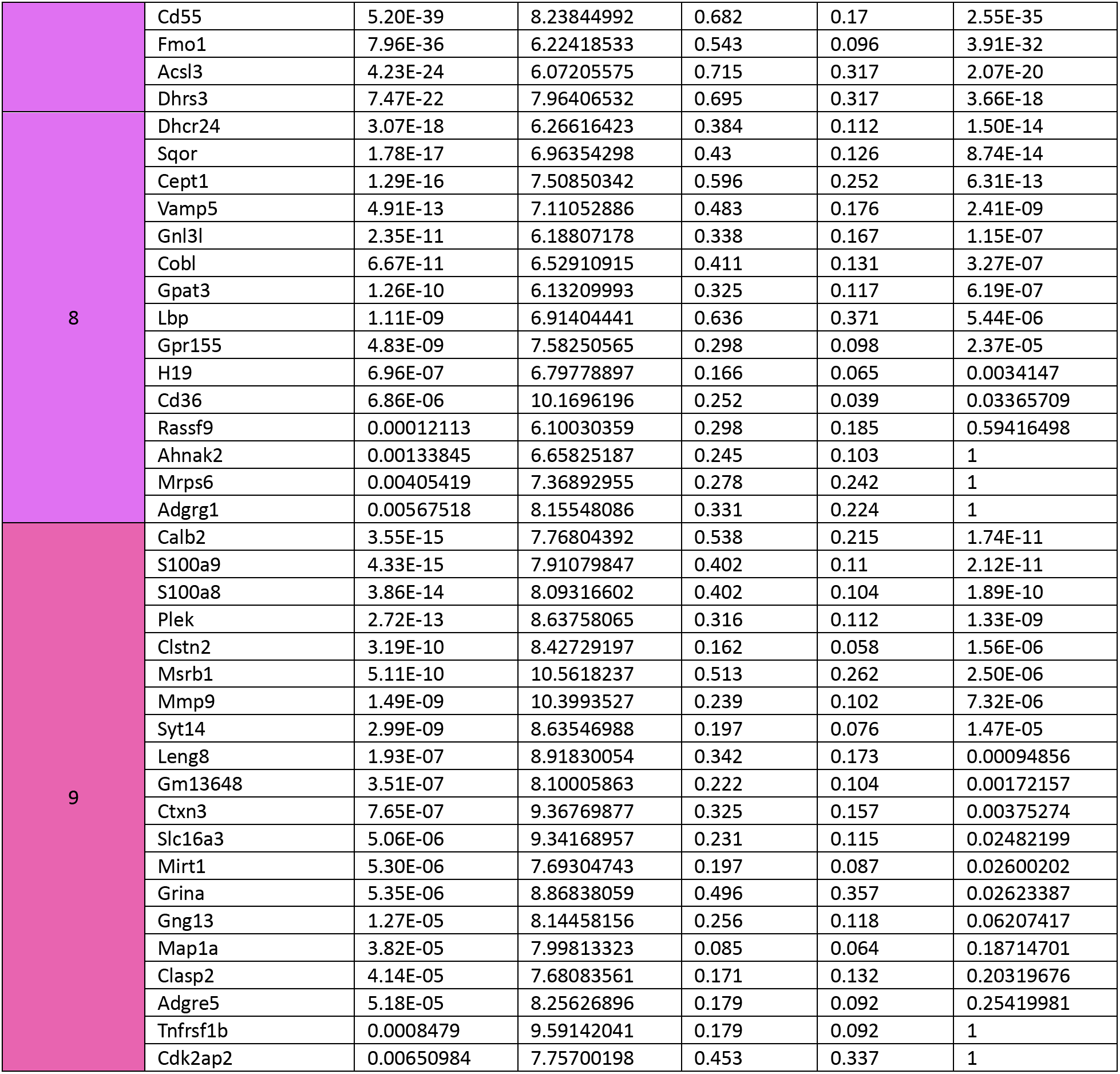
Top 20 Enriched Genes in each of the NSE Clusters.

